# Urban warming inverse contribution on risk of dengue transmission in the southeastern North America

**DOI:** 10.1101/2020.01.15.908020

**Authors:** Lorena M. Simon, Jesús N. Pinto-Ledezma, Robert R. Dunn, Thiago Rangel

## Abstract

1. Preventing diseases from becoming a problem where they are not is a common ground for disease ecology. The expectation for vector-borne diseases, especially those transmitted by mosquitos, is that warm and wet conditions favor vector traits increasing transmission potential. The advent of urbanization altering inner climate conditions hazards to increase mosquito’s transmission potential on “disease-free” cooler areas as a consequence of a warming urban heat island (UHI) effect.
2. We assessed the realism of the anticipated dengue transmission potential into the southern United States in a causal pathway with the ongoing UHI effect, vectors’ spatial distribution patterns, and exogenous environment; We also measured the climatic niche similarity between both dengue vectors species.
3. Our path model revealed that the UHI effect presents negative or no relation with dengue transmission potential. Instead, the surrounding non-urban temperature was rather suitable for the expected mosquitos’ transmission potential.
4. Both dengue vectors’ occurrence revealed to be more aggregated then expected by chance. These mosquitos’ density patterns were responsive to the warming effect of UHI-especially *Aedes Aegypti*-but not a reliable predictor for the anticipated dengue transmission potential pattern. The climatic niches of both vectors are not equivalent. Although currently highly overlapped, there is a wide space of their climatic niche still to be filled.
5. *Policy implications.* We highlight that the warming UHI effect on urban sites is not congruent with the expected suitability for dengue transmission. Instead, non-urban areas would be a better focus for dengue hazards into the southern United States. Our study also highlights the need for including low scale temperature on further mosquito-borne disease transmission models and track vectors niche filling under anthropogenic changes.

## 1. INTRODUCTION

Vector-borne pathogens are characterized by their dependence on vectors, in general arthropods (*e.g.*, mosquitoes), that feed on blood to proceed the infection cycle (Gubler, 2002). Their resulting diseases represent one of the greatest challenges faced by public health worldwide (World Health Organization, 2014). Some critical vector-borne diseases, such as; dengue, chikungunya, and Zika, which were formerly restricted to tropical and subtropical regions, have begun to spread into new parts of the world as a consequence of accidental introductions of vectors and pathogens along with changes in climate and habitat distributions (Gubler, 2001; Murray et al., 2015).

Despite the complex nature of vector-borne diseases transmission, understanding main drivers of its geographic spread is crucial for monitoring potential impacts on public health (World Health Organization, 2014). Vector transmission is linked with traits such as the biting rate, life span, and inner incubation period albeit also the abundance of vector species (Watts et al., 2018). Recently, studies have considered the temperature role on vector traits to assess the environmental suitability range for transmission capacity (*e.g.*, Brady et al., 2014; Ryan et al., 2019). This geographical perspective highlights the potential of macroecological analyses on disease ecology and public health strategies against the burden of disease transmission (Stephens et al., 2016). In this sense, the usage of transmission potential spatial patterns on causal structures with acknowledged exogenous drivers (*e.g.*, urban features), rise as a promising research area to infer causal pathways on disease ecology, ultimately orienting effective strategies on public health surveillance (Kraemer et al., 2019, Mordecai et al., 2019).

Some of the most consequential vectors species, including *Aedes aegypti* and *Aedes albopictus*, have lifestyles adapted to the ecology of urban settings and inner climatic conditions have the potential to favor their vector traits (Arnfield, 2003; Gloria-Soria et al., 2018). Urban sites might exhibit higher temperatures than surrounding, a phenomenon called ‘Urban Heat Island’ (UHI), and changes in the global climate and human population growth are expected to intensify the UHI conditions (Zhao et al., 2014; Manoli et al., 2019). As ectotherms, mosquito behavior, abundance, fitness and distribution patterns can be strongly affected by small changes in temperature (Amarasekare & Savage, 2011; Huey et al., 2012). In cooler regions, relative to mosquito species thermal optima, it is expected that species’ abundance might increase with UHI effect, particularly at range margins of mosquito species (Ladeau et al., 2015; Kraemer et al., 2019). In addition, even where mosquito species do not increase in abundance, their vectorial capacity might increase with the UHI effects (Araujo et al., 2015; Murdock et al., 2017). Conversely, UHI effects in areas already near the thermal maxima of a mosquito species may lead to decreases in their vectorial capacity (Mordecai et al., 2019). For instance, known upper thermal bounds for dengue transmission is 34.0 C° for *Ae. aegypti* and 29.4 C° for *Ae. albopictus* (Ryan et al., 2019). However, even though UHI effects might increase the potential for a disease outbreaks at the range margins of vector mosquitoes, UHI effects have received little attention in the infectious disease ecology (Misslin et al., 2016).

Dengue is a neglected disease that has rapidly expanded geographically over the last decades (Gubler, 2002; Ramos-Castañeda et al., 2017), and although *Ae. aegypti* was historically considered the main responsible for dengue urban transmission, *Ae. albopictus* has starred in recent major outbreak events (Lambrechts et al., 2010). Despite of their differing invasion timing and native origins (Kaplan et al., 2010), both vector species are currently listed among worst invasive organisms (Global Invasive Species Database, IUCN). In a recent future, both species are expected to spread farther north and south into temperate regions along with climate changes (Kraemer et al., 2019). Besides, the ongoing co-occurrence and continuing spread of both vectors in “dengue-free” areas, such as the southeastern North America, aggravates the temperature suitability predictions of dengue transmission into these areas under current and future climate conditions (Brady et al. 2014; Rosenberg et al., 2018; Messina et al., 2019), given the lack of heard immunization (Johnson et al., 2017).

In the present work, we aim to evaluate the realism of the geographical pattern on dengue transmission potential in the face of the urban heat island effect (UHI) and existing vectors distribution in the southeastern United States. To achieve our goal, we take two steps. Foremost, we build a correlative path structure (Fig. 1) to comprise the weight of UHI effect and other urban features on observed dengue potential pattern; and, as burden of dengue transmission largely reflects the distribution and density of the mosquito vectors, we also consider both the effect of urban features in increasing *Ae.* aegypti and *Ae. albopictus* clustering and the dengue transmission risk resulting from their distribution pattern. Second, as the niche similarity between these important mosquito vectors is unclear, we additionally use a niche overlap approach to compare the climatic niches of both species assuming that, in spite of sharing similar geographical spaces, they do not have equivalent niches, which would result in low overlap between vectors niche and in current avoidance of dengue outbreak on the region.

**Figure 1.**
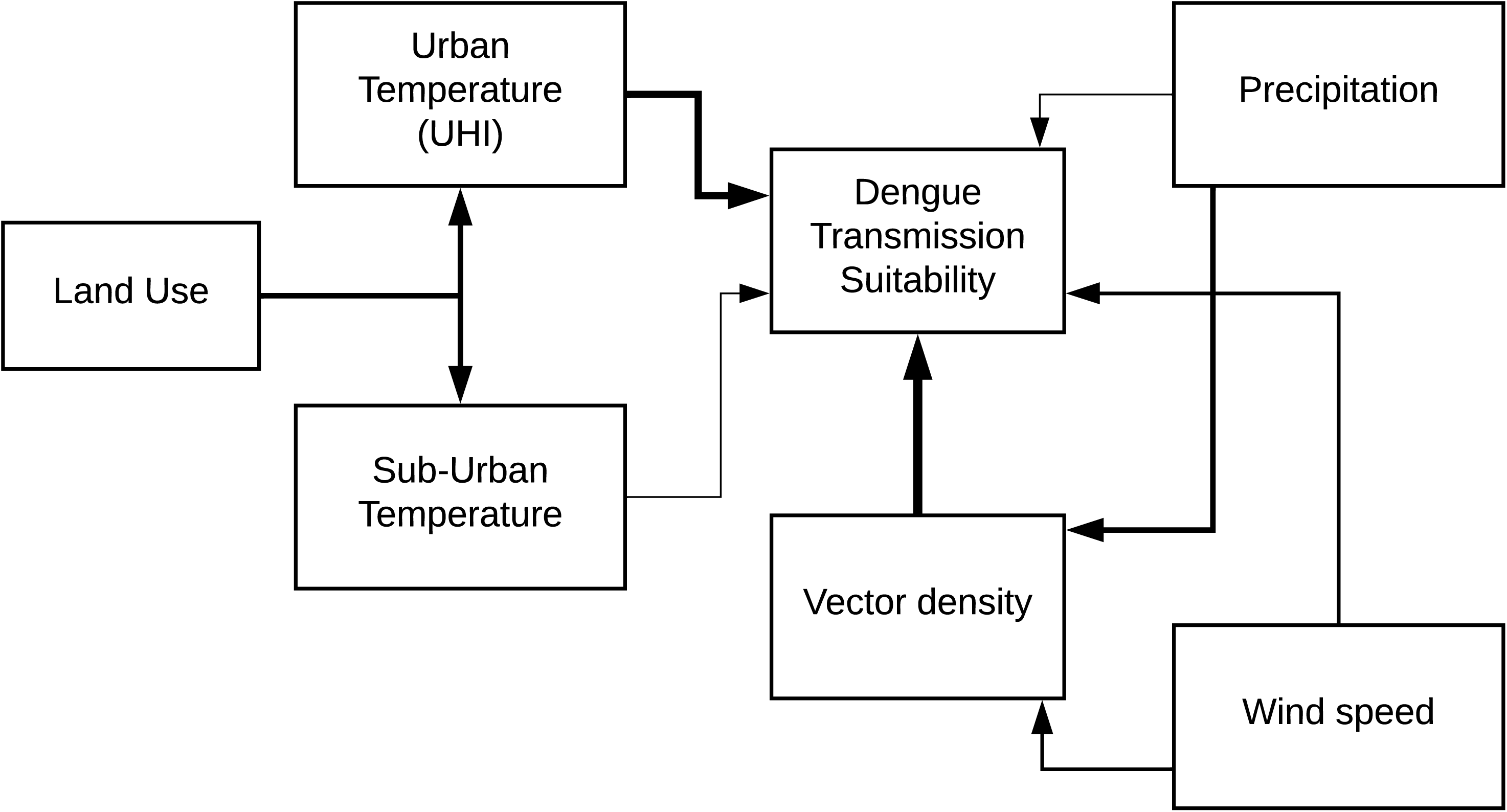
Conceptual Path model defining the expected relations between predictors and response.

## 2. MATERIALS AND METHODS

### (a) Data

#### (i) Vector species occurrence

The *Ae. aegypti* (*Stegomyia aegypti*) and *Ae. albopictus* (*Stegomyia albopicta*) occurrence was obtained from Kraemer et al. (2015) available at http://datadryad.org/resource/doi:10.5061/dryad.47v3c and improved with last published records (Johnson et al., 2017). This comprehensive dataset is a compilation of occurrence point records over the last 57 years (1960 – 2017) documented in previous studies– Kraemer et al., 2015; Hahn et al. 2016; 2017. To latter evaluate each specie density in southeastern United States we selected the occurrence points located into the region, ultimately comprising 1227 and 217 records of *Ae. albopictus* and *Ae. aegypti*, respectively.

#### (ii) Dengue transmission suitability

To assess the geographic range of dengue transmission potential we used Brady’s *et al.* (2014) global consensus map of vector transmission suitability based on temperature, from which we extracted the information within the southeastern United States. This map is a result of a mechanistic model derived from experimental data that assess vector traits of dengue transmission (*e.g.*, mosquito survival; extrinsic incubation period [EIP]) based on temperature effect, separately for *Ae. aegypti* and *Ae. albopictus*, and spatialized using a global temperature dataset with 1km resolution (see Brady *et al.* (2014) for modeling approach details). Their output predicted areas where global temperature support year-round dengue transmission given the presence of an infected individual (*i.e.* the basic reproduction number, R0) ranging from 0 to 1, for each vector species, where within pixel values closest to 1 indicate a higher potential for the virus transmission.

#### (iii) Meteorology and land use

To test the effect of urban differential temperature on dengue transmission suitability we used the UHI dataset from ‘NASA Socioeconomic Data and Applications Center’ (SEDAC, 2016), available at http://sedac.ciesin.columbia.edu/data/set/sdei-global-uhi-2013/data-download. The UHI data comprises the estimate of summer daytime maximum and nighttime minimum surface temperature within urban extent and surrounding non-urban areas (buffer of 10 km), and the difference between them, in Celsius degrees. Here we used both daytime and nighttime temperatures, once vector activity is referred to be even superior at nighttime than it is in daylight (Stoddard et al., 2009). The global GeoTIFF is in the resolution of 30 arc-seconds (∼1Km), on which we made a subset based on southeastern United States area.

To account for the influence of other urban-modified features on vectors density and dengue transmission we obtained the data referent to precipitation and wind speed from NASA Langley Research Center (LaRC) POWER Project, available at https://power.larc.nasa.gov/data-access-viewer/. Both variables represent the average annual information in a 0.5° global grid. The wind speed data is scaled on 2 meters elevation, accounting for the limited space of mosquito’s activity (Reisen et al., 2003; Guerra et al., 2014). The land-cover features, known to increase mosquito vectors density due to anthropogenic changes favoring species associated with urban areas (Beaulieu et al., 2019), were obtained in 1-km resolution (Tuanmu & Jetz, 2014). The data account for 7 land-cover classes (*i.e.*, Evergreen/Deciduous Needleleaf Trees, Deciduous Broadleaf Trees, Mixed/Other Trees, Herbaceous vegetation, Cultivated and Managed Vegetation, Regularly Flooded Vegetation and Urban/Built-up) chosen based on their matter on vector settlement and ultimate dengue transmission (Guerra et al., 2014).

Finally, to build and later compare both vector species niche we delimitated the niche boundaries using bioclimatic variables, which are widely accepted given its robustness to represent seasonal trends and physiological constrains of species (Lobo et al., 2010). In this sense, we used all 19 bioclimatic variables from WorldClim dataset on the resolution of 30 arc-seconds, available at http://www.worldclim.org/.

### (b) Analysis

#### (i) Density estimation

For the estimation of vector density across southeastern U.S., we used the point pattern approach based on species occurrence data. Firstly, to account for bias in point density estimation, we applied the rarefaction curve, commonly used to quantify bias in presence counting measures (Gotelli & Colwell, 2001), with the R package *iNEXT* (Hsieh et al., 2019). Then, we used the R package *spatstat* (Baddeley et al., 2015), where we performed a near neighbor analyses (ANN) between vector occurrence records and compared with a commonly used null model based on the distribution of simulated ANN values given the Complete Spatial Random (CSR) point process (Wiegand & Moloney, 2004; Baddeley et al., 2014). To generate a vector density raster, we used the Kernel density estimation to interpolate around each point, for which we established the bandwidth value of 0.5 to give weight to distant points contribution on density estimation using the R package *KernSmooth* (Wand, 2015).

#### (ii) Path model

To recover the underlying direct and indirect causal mechanisms between UHI and other features on dengue transmission suitability on the southeastern United States (Fig. 1) we used the structural equation modeling (SEM) approach with the R package *lavaan* (Rosseel, 2012). Usually, the SEM model is applied to mediate causal assumptions, which assumes that the presumed explanatory variables can influence an outcome directly and indirectly through other variables (Fan et al., 2016). However, SEM does not account for spatial information and the autocorrelation that frequently arise when dealing with spatially explicit structures, which ultimately inflates the type I error given the lack of independence between observations across space (Legendre & Legendre, 1998).

Aiming to consider the spatial autocorrelation and provide unbiased regression coefficients we used eigenvector-based spatial filters, which consist on extracting the eigenvectors of a distance matrix describing the spatial structure of the data and adding them as additional predictors into the SEM model (Griffith, 2003). First, we extracted the geographical coordinates along southeastern U.S. to build a distance matrix, which was truncated at the distance of 300 km based on a previous evaluation of the Moran’s I correlogram. Then, the truncated matrix was submitted to a principal coordinate analyses (PCO) and its resultant eigenvectors were selected as predictors based on significance of each partial regression coefficients (following Borcard & Legendre, 2002). For the spatial filter approach, we used the R packages *letsR* (Vilela & Villalobos, 2015) and *ecodist* (Goslee et al., 2007.

#### (iii) Niche overlap

In order to estimate the overlap between dengue vectors niches and test the hypothesis of niche non-equivalence we used the framework proposed by Broennimann et al. (2012). The method assesses niche overlap by calibrating a principal component analysis on the environmental space (PCA-env) and use kernel density smoothing to correct potential sampling bias (Broennimann et al., 2012). We used the R package *ecospat* (Di Cola et al., 2017) to pull the bioclimatic information, according to the species occurrence, and create a background environmental space to perform the PCA. Thus, the 1^st^ and 2^nd^ output axes were used to create a 100 x 100 occurrence density grid representing each specie niche. The estimated niches was overlapped and the degree of intersection was assessed using Schoener’s D metric (following Warren et al. 2008)– which ranges between 1 (*i.e*., complete overlap) and 0 (*i.e.*, no overlap) –and compared with 100 random simulated overlap index distribution to test for niche equivalence and niche similarity.

## 3. RESULTS

### (a) Density estimation

The test for vector occurrence point pattern clustering/dispersion on the southeastern region of United States showed that, when compared with a null distribution of average distance among geographic points, the distance between occurrence points density, for both *Ae. albopictus* and *Ae. aegypti*, are greater than expected. Even though, *Ae. aegypti* density pattern revealed higher concentration into Florida and Louisiana, while *Ae. albopictus* showed a more diffused occurrence density, with higher weight into the northern portion of southeastern U.S. like Virginia (Fig. 4).

### (b) Path model

The SEM revealed that the combined influence of diurnal and nocturnal UHI, wind speed, precipitation and land-use-including spatial filters - explained respectively 49% and 54% of the variance on *Ae. aegypti* and *Ae. albopictus* density (Fig. 2). In addition, the interaction between vectors occurrence density, UHI and further predictors explained 92% and 90% of variation on dengue transmission suitability respectively by *Ae. aegypti* and *Ae. albopictus* into the southeastern U.S. (Fig. 2). The addition of spatial filters on SEM structure to take in account the unknown endogenous and exogenous influence shaping dengue transmission suitability pattern, improved the model fit based on Akaike information criterion (AIC) and r square. The inclusion of 20 spatial filters on *Ae. aegypti* path model adjusted the AIC from 15356.419 to 11631.864 and the R^2^ from 0.5 to 0.9, and the 30 spatial filters included on *Ae. albopictus* model adjusted the AIC from 15655.011 to 11845.485 and the R^2^ from 0.34 to 0.9 (Table 1).

**Table 1.** Table showing the outcome correlation coefficients of each path resulting from SEM analyses and the respective adjust of each model given the consideration autocorrelation by spatial filters.

|  | Vector: *Ae. Aegypti* |  | Vector: *Ae. albopictus* |  |
| --- | --- | --- | --- | --- |
|  | Parameter estimate | Standard Error | Parameter estimate | Standard Error |
| **Dengue transmission suitability ~** |  |  |  |  |
| Mosquito Density | -0.02* | 0.01 | -0.01 | 0.01 |
| UHI Daytime | -0.03*** | 0.02 | -0.03*** | 0.02 |
| UHI Nighttime | 0.001 | 0.01 | 0.002 | 0.02 |
| Wind Speed | 0.07*** | 0.01 | 0.013*** | 0.03 |
| Precipitation | -0.03 | 0.04 | -0.08 | 0.04 |
| Spatial Filters | 0.04* | 0.01 | 0.02*** | 0.01 |
| **Mosquito density ~** |  |  |  |  |
| UHI Daytime | 0.05*** | 0.03 | 0.02 | 0.02 |
| UHI Nighttime | -0.04*** | 0.02 | -0.04* | 0.01 |
| Wind Speed | -0.19*** | 0.03 | -0.07*** | 0.01 |
| Precipitation | 0.31*** | 0.06 | 0.23*** | 0.06 |
| Spatial Filters | 0.04 | 0.04 | 0.03* | 0.02 |
| AIC | 11631.864 |  | 11845.485 |  |
| AIC (nf) | 15356.419 |  | 15655.011 |  |

\*p≤0.10.
\*\*\*p≤0.01.
(nf) No spatial Filter.

**Figure 2.**
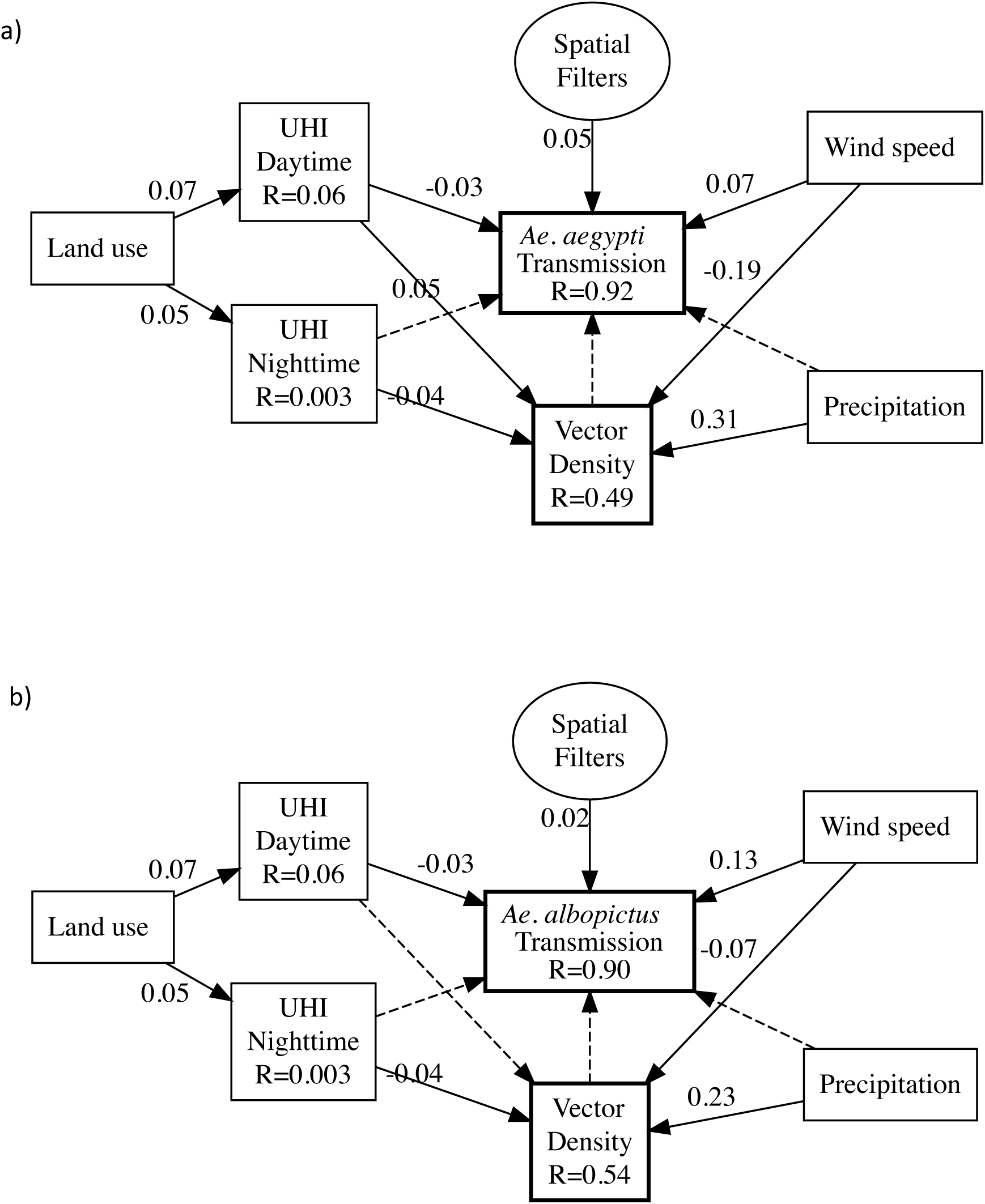
Structural equation model outcome for a) *Ae. albopictus* and b) *Ae. aegypti*. R^2^ is shown for de dependent variable. The values associated with arrows are standardized regression coefficients and dashed arrow indicate non-significance path coefficient (P > 0,05).

The resulting SEM causal path indicated that daytime UHI (Fig. S1) is negatively correlated with dengue transmission suitability (β = −0.03; sites with a greater UHI effect are less suitable for transmission by vectors) but positive with the density of both vectors (β = 0.05; 0.02; sites with a greater UHI effect have more of both mosquito species), although the effect on the density pattern of *Ae. albopictus* is not significant. In contrast to the effect of daytime UHI, the UHI effect during nighttime (Fig. S1) was not significantly correlated with dengue transmission suitability in the southeastern U.S. Interestingly, the density of both vector species was negatively correlated with nighttime UHI (β = −0.04) (Fig. 2; Table 1), such that the effect of daytime UHI and nighttime UHI were in the opposite directions for the mosquito species agglomeration pattern. Precipitation was strongly positively correlated with the density of both vectors (β = 0.31; 0.23), while its effect in dengue transmission suitability was not significant. The SEM path output also indicated that wind speed has a high negative effect on the density of both vector species (β = −0.19; −0.07), however it showed a positive association with dengue transmission suitability by *Ae. aegypti* (β = 0.07) and *Ae. albopictus* (β = 0.013). The effect of land-use over dengue predictors was indirect via its influence on UHI (β = 0.07; 0.05). Moreover, the occurrence density of the two vectors was not correlated with dengue transmission suitability in the southeastern U.S. (Fig. 2; Table 1).

### (c) Niche overlap

Schoener’s D niche overlap index revealed a high level of overlap between Ae. *albopictus* and *Ae. aegypti* niches (Fig. 3). The niche similarity test showed that niche overlap comparisons between one randomly distributed over the unchanged other (1 -> 2) and vice versa (2 -> 1) had a Schoener’s D of 0.44, thus distant from a completely unrelated scenario (*i.e.*, D = 0). The vector species niches are represented by the 1^st^ axes of the PCA-env, that is associated with temperature-related bioclimatic variables, and by the 2^nd^ axes, that is associated with precipitation-related variables. In spite of the high niche overlap and similarity of both vectors species, the result of the one-tailed niche equivalence test showed a significantly lower niche equivalence between both main dengue vectors (p-value = 0.001).

**Figure 3.**
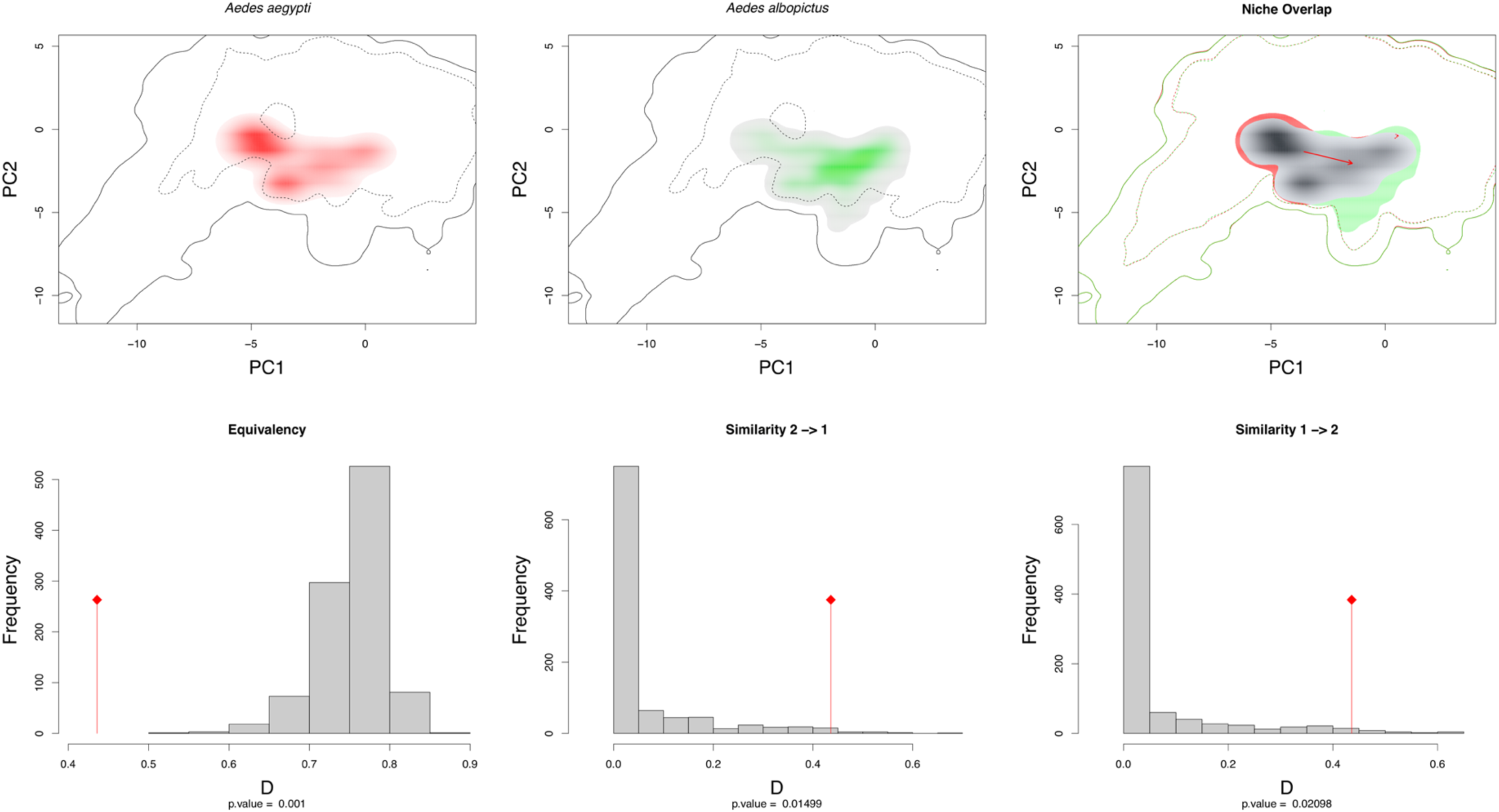
Niche overlap and niche equivalence result showing the occupied climatic niches of *Aedes albopictus* and *Aedes aegypti* (Top left) in the niche space available, and the amount of niche overlap between them (Top right). Both ways intersection (Bottom right) showed higher overlap degree than expected by the null model (*i.e.*, Schoener’s D > 0.4). In spite of highly overlapped, their niche spaces are not equal (Bottom left).

**Figure 4.**
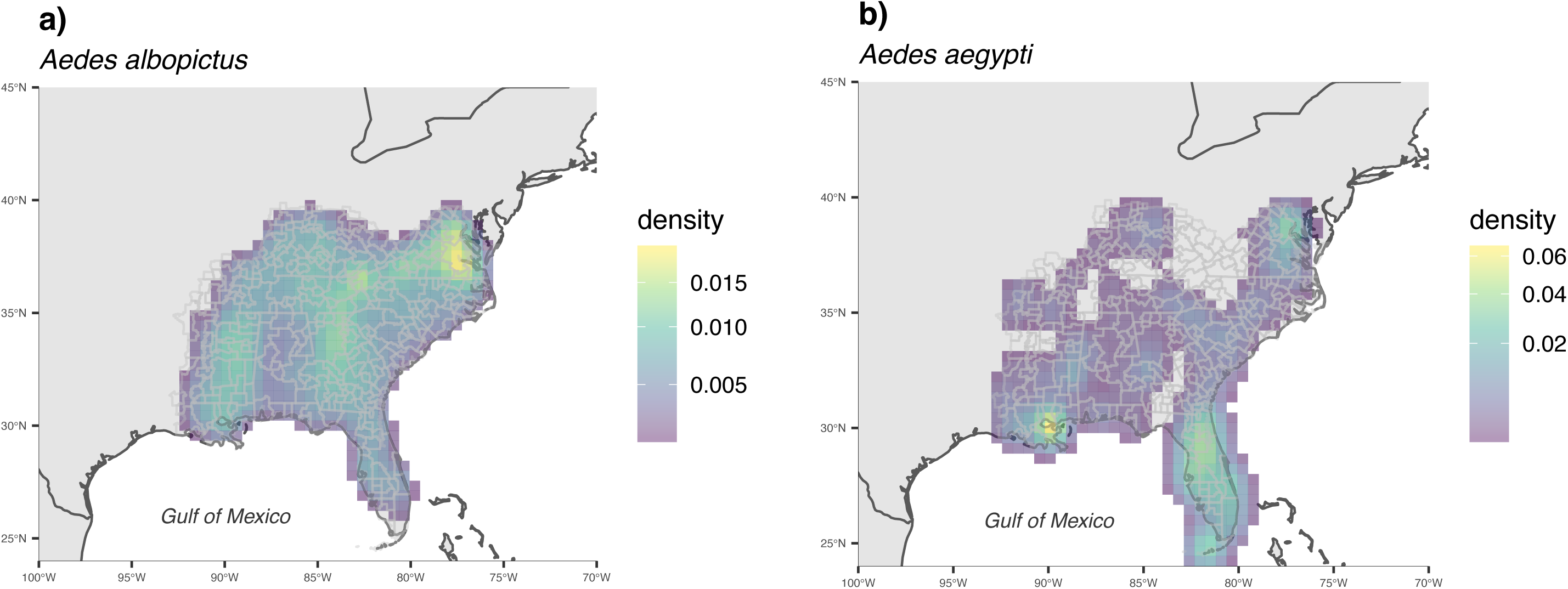
*Aedes albopictus* (a) and *Aedes aegypti* (b) spatial density estimation based on Kernel smooth approach of available occurrence records.

## 4. DISCUSSION

Contrary to previous expectations that urban heat islands (UHI) effect might favor dengue transmission, our results suggest that areas under UHI stress have lower risk of the disease spread by its both main vectors. The threat of dengue transmission potential into the southeastern U.S. has been placed decades ago based on mosquito vectors invasion (Monath, 1994). The incorporation of temperature on disease transmission models later reinforced the predictions of *Ae. aegypti* and *Ae. albopictus* potential dengue transmission into the region under current and future global climate (Brady et al., 2014; Messina et al., 2019). However, here we show that the downscaled temperature difference between warmer urban and cooler sub-urban areas– the so called UHI –have a contrasting inverse relation with dengue transmission potential (Fig. 2; Table 1). Contrary to the expectation that warmer conditions generally promote mosquito borne disease (Morin et al. 2015; Thomson et al. 2017). Still, UHI effect, wind speed, and precipitation all together were highly congruent with the expectation of dengue transmission suitability on the southeastern U.S., which aligns with the concern of urbanization style shaping the probability of mosquito-borne disease transmission (Gubler, 2011).

The starting point for realized dengue transmission depends primarily on the presence of virus strains, susceptible host population and competent vectors (Gubler, 2011). Year-round adequate temperature determining transmission competence of mosquito vectors (Ryan et al., 2019), in combination with precipitation, ultimately zenith seasonal dengue cycles into tropical endemic areas (Van Panhuis et al., 2015). In contrast, cooler subtropics are expected to avoid transmission following an unimodal variation on vector-borne transmission that limits dengue around 18 C° (Ladeau, 2015; Mordecai et al., 2019). Here we outline that on summer the combination of temperature with other environmental features is congruent with dengue transmission potential and vectors accumulation on the subtropical southeastern U.S. (Fig. 2; Table 1). However, contrary to the expected higher dengue probability in warmer urban temperature (Halstead, 2008), transmission potential showed conformity with the sub-urban lower temperature. In fact, previous works highlighted that mosquito transmission increased around lower temperature ranges (Carrington et al., 2013), advocating for a thermoregulation scape from urban to bordering greener sites where environment afford thermal respite (Huey, 2012; Misslin et al. 2016). Still, the dengue potential contrast we found between urban and surrounding areas might also reflect the scale considered to predict the temperature range of dengue transmission potential, which ignores low scale environment where transmission takes place.

In cities where dengue transmission is a seasonal event, human population cluster share space with high density of mosquitoes, usually a strong predictor for arboviral transmission potential (Halstead, 2008; Ladeau et al., 2015). Our results otherwise showed that the density derived from occurrence data was not a good predictor of the dengue potential predicted by the temperature-based transmission model in the southeastern U.S. This result might either represent that the mosquito records within the studied area are not sufficient to predict the emergence of dengue, or that both species aggregation does not overlap the dengue transmission suitability areas. Although previous works have indicated a positive association between vector density and disease incidence (Walk et al., 2009), this association does not occur in all cases (Halstead, 2008). For instance, in Singapore the extreme reduction of *Ae. aegypti* density did not avoid the continued dengue infection (Chan, 1985). Here we found that both mosquito species densities were more related with urban then suburban temperatures (Table S1), congruent with urban microclimates favoring vector population growth-while it has not reached thermal performance peak -(Huey et al., 2012; Mordecai et al., 2019). Additionally, day and nighttime UHI range revealed fully relation with *Ae. aegypti* density (Figs. 2, S1), a primarily urban specie when compared with the *Ae. albopictus*, which dominates in suburban areas (Beaulieu et al., 2019).

The southeastern U.S. have a particular precipitation regime with much higher humidity than other U.S. locations. Consequently, the UHI effect-which follows precipitation gradient (Manoli et al., 2019) -is increased in this region, where annual UHI effect is around 3.9 C° higher than dryer U.S. regions (Zhao et al., 2014). Besides the indirect effect on urban temperature higher precipitation is also expected to increase mosquito density by increasing breeding sites and oviposition (Halstead, 2008), and our results showed a positive association between precipitation and both species’ densities, supporting this prediction. However, precipitation was not a good support for the dengue suitability range expected by the global temperature model. In this sense, the background effect of higher precipitation on the southeastern U.S. UHI might indirectly impose thermal limitations to dengue transmission range, even where wind speed is expected to facilitate the transmission contact (Cummins et al., 2012). To fully comprehend this complex association between precipitation and UHI on dengue transmission, further works should include the urban differential climate into mosquito-borne disease transmission models.

The niche comparison revealed that, in spite of distinct invasion time by dengue vectors into the southeastern U.S.-where there is a niche conservatism evidence for both species (Cunze et al., 2018) -their occupied climatic space markedly overlap (Fig. 4). Accordingly, given the observed low prediction of dengue transmission suitability by vector density on the southeastern U.S., the ongoing climatic space occupied by both species may be the main factor avoiding a dengue outbreak into the region, when other important features are favorable (*e.g.*, autochthonous virus, lack of heard immunity). However, our results also highlight that the climatic niche space of both species are not completely fulfilled (solid line Fig. 3), which ultimately indicate future potential for dengue transmission due to range expansion. In fact, Kraemer et al. (2019) showed that there is strong evidence for future *Ae. aegypti* and *Ae. albopictus* range expansion poleward with anthropogenic pressure, ultimately fulfilling the remaining suitable climatic space and increasing the risk of dengue transmission into subtropical areas like the southeastern U.S.

## 5. CONCLUSIONS

Here we highlight that the southeastern U.S suburban areas show higher realism with expected dengue transmission thermal bounds. If the expected dengue potential range is accurate, suburban great transmission suitability ultimately represent higher risk of infectious contact between humans and competent vectors once this is a residential zone. However, vectors density pattern did not show correspondence with the suitability range based on global temperature, which might indicate an underestimation of dengue risk on warmer urban areas. In addition, the niche space currently occupied by both vectors are similar but not equivalent, and part of their climatic niche remain unfiled, representing an ahead risk of vectors population grow into areas of dengue transmission competence. In this sense, range expansion of both species under anthropogenic and climatic changes claim that the combat of mosquitoes must be intersected in areas where the contact of hosts and competent vectors represents a risk. Accordingly, here we suggest that considering the UHI effect on further predictive dengue transmission models might be crucial to accurately identify areas of dengue transmission risk. Moreover, further research is still needed to address the heat suitability on mosquito’s traits to transmit other concerning viruses such as Zika and chikungunya (Carlson et al., 2018), in the light of virus and vector coevolution and evolutionary adaptation to new environments.

## DATA ACCESSIBILITY

This article has no additional data.

## AUTHORS’ CONTRIBUTIONS

L.M.S. and R.R.D. conceived the project; L.M.S managed the project; L.M.S and J.N.P-L. conducted the analyses; all authors contributed to the project and/or drafting of the manuscript.

## COMPETING INTERESTS

The authors declare that the research was conducted in the absence of any commercial or financial relationships that could be construed as a potential conflict of interest.

## FUNDING

L.M.S. was supported by CAPES fellowship over PROEX PhD and the Sandwich Doctoral Program Abroad-PDSE. J.N.P-L. was supported by the University of Minnesota College of Biological Sciences’ Grand Challenges in Biology Postdoctoral Program.

## ACKNOWLEDGMENTS

We thank Brian Wiegmann and John Soghigian for the rich discussions about mosquito ecology and behavior.

