## Supplementary figures and images for "Urban warming inverse contribution on risk of dengue transmission in the southeastern North America"

### Supplemental Fig. S1

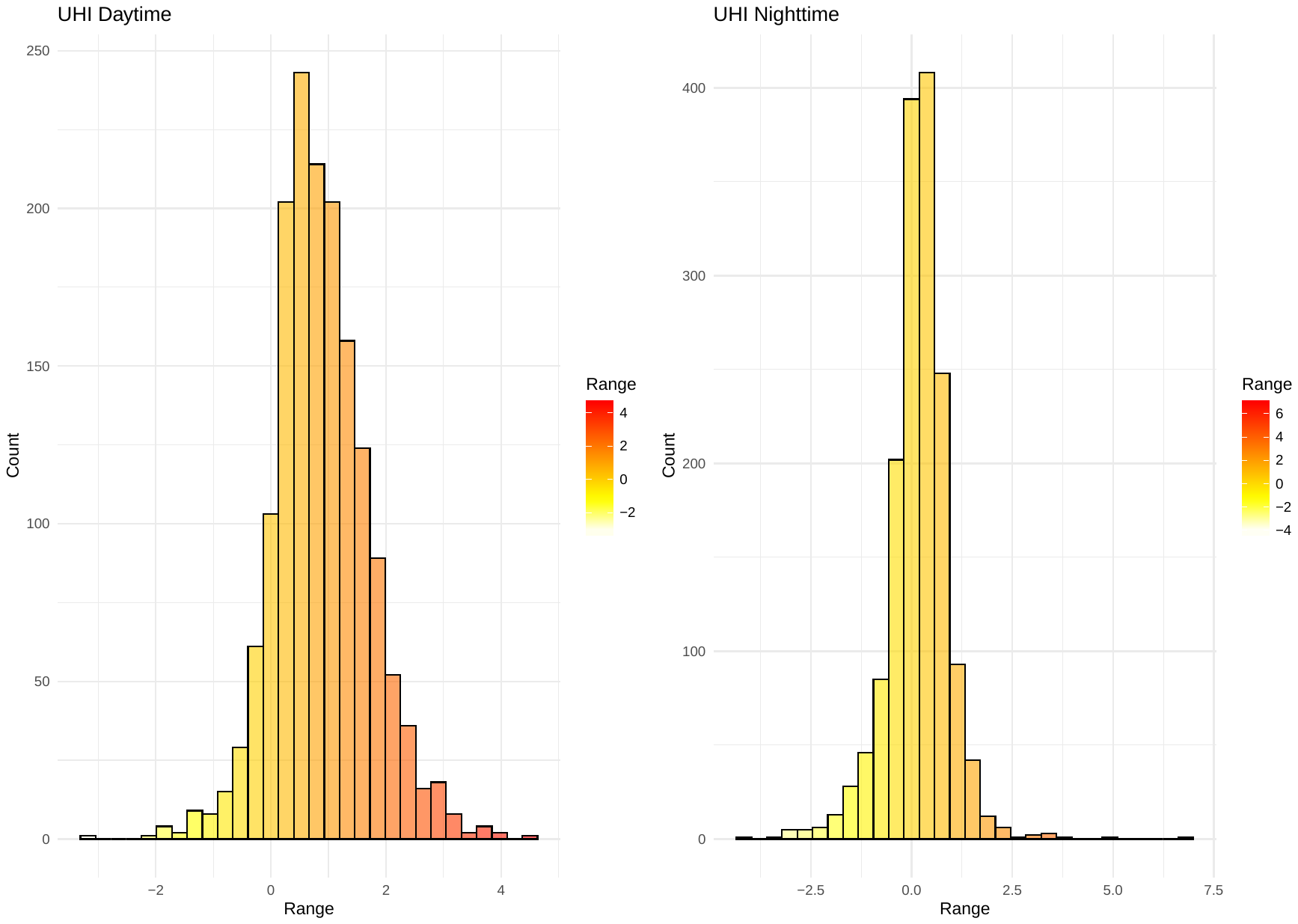

**Figure S1:** UHI daytime data range (left) and UHI nighttime data range (right).
