## Supplemental Table S1 for "Urban warming inverse contribution on risk of dengue transmission in the southeastern North America"

| **Table S1** Table showing the outcome from further SEM analyses construction on which we considered urban and sub-urban temperatures separately. | | | | | |
| --- | --- | --- | --- | --- | --- |
|  | Vector: *Ae. aegypti* | |  | Vector: *Ae. albopictus* | |
|  | **Parameter estimate** | **Standard Error** |  | **Parameter estimate** | **Standard Error** |
| **Dengue transmission suitability ~** |  |  |  |  |  |
| Mosquito Density | -0.01* | 0.01 |  | 0.02* | 0.01 |
| Urban Daytime Temperature | -0.04*** | 0.01 |  | -0.05*** | 0.01 |
| Sub-urban Daytime Temperature | 0.18*** | 0.01 |  | 0.12*** | 0.01 |
| Urban Nighttime Temperature | 0.02*** | 0.02 |  | 0.02*** | 0.01 |
| Sub-urban Nighttime Temperature | 0.45*** | 0.01 |  | 0.59*** | 0.01 |
| Wind Speed | -0.01 | 0.01 |  | 0.05*** | 0.01 |
| Precipitation | -0.15*** | 0.03 |  | -0.26*** | 0.04 |
| Spatial Filters | 0.03*** | 0.01 |  | 0.51*** | 0.01 |
| **Mosquito density ~** |  |  |  |  |  |
| Urban Daytime Temperature | 0.15*** | 0.03 |  | 0.07*** | 0.01 |
| Sub-urban Daytime Temperature | -0.02 | 0.02 |  | 0.09*** | 0.02 |
| Urban Nighttime Temperature | -0.16*** | 0.04 |  | -0.12*** | 0.02 |
| Sub-urban Nighttime Temperature | 0.05*** | 0.02 |  | -0.08*** | 0.02 |
| Wind Speed | -0.22*** | 0.03 |  | -0.11*** | 0.03 |
| Precipitation | 0.28*** | 0.05 |  | 0.21*** | 0.04 |
| Spatial Filters | 0.21* | 0.02 |  | 0.76* | 0.02 |
| ML | 12992.465 |  |  | 13313.190 |  |
| AIC | 21013.03 |  |  | 21072.68 |  |
| AIC (nf) | 23654.16 |  |  | 23843.9 |  |
| *p≤0.10. |  |  |  |  |  |
| ***p≤0.01. |  |  |  |  |  |
| (nf) No spatial Filter. |  |  |  |  |  |
